# Efficient Multi-Label Attribute Classification and Recognition of Microbiological Bacteria Based on Deep Learning and model fine-tuning

**DOI:** 10.1101/2022.10.05.511056

**Authors:** Duidi Wu, Haiqing Huang, Shuo Zhang, Jin Qi, Dong Wang, Jie Hu

## Abstract

Bacterial vaginosis (BV) is the most common gynecological complaint affecting health of a large percentage of women worldwide. Traditional manual microscopy methods are expensive and time-consuming, to improve accuracy and efficiency, automated bacterial identification devices with detection intelligence algorithms are urgently needed. We propose a Fine-tuned SmallerVGG (FTS-VGG) deep convolutional network model based multi-label classification method for bacteria. Comparison experiments were deployed on several advanced backbone networks, including transfer learning on pre-trained VGG19, demonstrating that the proposed method achieves the advantages of being lighter, faster, more accurate and more efficient. Due to the high cost of time and expertise of experienced clinicians, we use random erasing for data augmentation to address the challenge of dataset collection and annotation, experiments demonstrate its robustness to occlusion. The proposed method has theoretical and practical implications, as well as the potential to be widely extended to other microscopic imaging applications.

## 1 INTRODUCTION

Bacterial vaginosis is one of the common vaginal infections in women of reproductive age. Domestic survey data show that bacterial vaginosis accounts for about 11% of women in health check-ups and 36%-60% of patients with vaginal inflammation in gynecological clinics. Complications associated with bacterial vaginosis are more frequent and can cause inflammatory pelvic diseases, post gynecological surgical infections and infertility. Bacterial vaginosis also increases the risk of infections such as Human Papilloma Virus (HPV), Human Immunodeficiency Virus (HIV) [1], Neisseria gonorrhoeae, Chlamydia trachomatis [2], Herpes simplex virus type 2 infections, etc [3][4].

The most common causes of vaginitis are bacterial vaginosis, vulvovaginal candidiasis, and trichomoniasis. Bacterial vaginosis is implicated in 40% to 50% of cases when a cause is identified. Noninfectious causes, including atrophic, irritant, allergic, and inflammatory vaginitis, are less common and account for 5% to 10% of vaginitis cases [5].

The diagnosis of bacterial vaginosis is currently based on the Amsel clinical criteria [6] and the Gram stain Nugent score [7] diagnostic criteria. Rapid detection of pathogenic microorganisms is of key importance to prevent potential diseases and safeguard health, safety and well-being. Vaginitis bacteria are not suitable for general pathogen detection methods such as culture colony count and polymerase chain reaction (PCR) [8][9]. Manual microscopic analysis of conventional bacterial assays may take up more days to yield results, and may have high false-positive results due to human errors, which cannot meet the needs of large number of samples examined. To improve efficiency and accuracy, computer-aided diagnostic methods based on deep learning are widely developed to automatically recognize medical images for disease detection [10].

## 2 RELATED WORK

### 2.1 Deep Learning BV Detection

With special multilayer neural networks and deep learning strategies, Convolutional Neural Networks (CNN) are able to recognize visual information directly from pixel images with minimal pre-processing, showing excellent performance in research such as image classification, semantic segmentation, and target detection, etc. Existing bacterial identification methods usually make architectural modifications on the basis of image classification, mostly follow the LeNet-5 [11], GoogLeNet [12], CaffeNet [13], AlexNet [14], VGG-16 [15] and ResNet50 [16] backbone.

Deep convolutional networks can outperform many traditional image processing and machine learning methods due to their high efficiency in extracting features from raw data. For research on diagnostic aids for medical images, general ideas mainly include image segmentation-based and target recognition-based detection. The small size, large number and splitting overlapping of bacteria make segmentation challenging. Song et al. [17] proposed a quantitative analysis of bacterial approach to diagnose BV, with a three-step automatic diagnosis method of region segmentation, overlapping division and bacterial morphological type classification. Wang et al. [18] used two convolutional neural networks with encoder-decoder architecture for trichomonas vaginalis detection, considered the motion characteristics of trichomonads and used *video detection* methods. The first network learn the difference between frames and utilizes the optical flow information to perform rough detection, then the second network corrects the coarse contours to perform fine detection. Wang et al.[19] optimized a deep CNN model to quantify Gram staining to achieve automatic classification of Nugent scores. Considering the loss of tiny cellular information due to compression of image resolution, they referred to EfficientNet [20] and developed *NugentNet* with two down-sample convolutional layers, scaled the network depth to adapt to the resolution and extract the detailed information about the bacteria.

In response to the difficulty of acquiring large amounts of labeled data for supervised models, some scholars have applied the ideas of *transfer learning* and *fine-tuning*. Hao.et al. [21] presented a data-efficient framework for the identification of vaginitis based on *transfer learning* and *active learning* strategies. Peng et al. [22] used fine-tuned InceptionResNetV2 for the automatic identification task of three vaginal diseases (BV, AV and VVC). It was showed that transfer learning can reduce the required manual labeling by roughly 73%.

However, a limitation of existing methods is that most studies only perform simple automatic binary classification of negatives and positives or add the identification of an intermediate state, without the ability to identify multiple bacterial forms simultaneously, which means new bacterial types requires retraining the corresponding network.

### 2.2 Contribution and Structure

In this work, we propose an efficient deep learning-based multi-label classification and identification method for vaginitis bacteria. The proposed method uses SmallerVGG convolutional networks as the backbone framework and performs model fine-tuning, apply multiple data augmentation methods to expand the native data under label consistency. The model trained by random erasure methods enhances the robustness of the network to occlusion conditions while reducing the cost of expensive labeling, which is expected to shorten the development cycle of medical devices for effective market application.

For the problem of identifying multiple bacterial properties simultaneously in common cases, there has not been much advanced work, which is the focus of this work, and the main contributions of this paper include:

- Establish an image dataset of vaginosis bacteria annotated with 9 category attributes named VB9, containing common bacteria such as trichomonads, spores, cocci, bacilli etc. and conventional cells such as white blood cells.
- Enhance raw data with Random Erasing Augmentation (REA), crop and rotation, etc., and verify their effectiveness and robustness under partial occlusion.
- Provide a fine-tuned SmallerVgg (FTS-VGG) deep learning-based method that implements multi-label classification for bacterial images, and demonstrate the advantages of the proposed model being faster, lighter and more accurate.

The rest of the paper is organized as follows: Section 3 introduces the dataset and several effective data enhancement methods, Section 4 describes the deployment of advanced deep learning models for bacterial multi-label classification, Section 5 describes the experimental part, mainly includes model testing, metric-based quantitative evaluation and comparison, Section 6 provides a discussion and conclusion.

## 3 DATASET

### 3.1 Source and Annotation

Our medical-industrial crossover team worked with the Shanghai Sixth People’s Hospital obtained representative microscopic images of vaginitis bacteria, which were screened by technicians with extensive clinical experience. We built a vaginal inflammation bacteria dataset containing 1560 microscopic images with a resolution of 1920 × 1200 pixels. In addition to conventional epithelial cells, the multi-category attributes include 9 cell types shown in Fig 1, covering common normal flora and inflammatory bacteria, namely clue cells, trichomonas, spores, cocci, naked nucleus cells, mycelium, white blood cells, germ tubes, and bacilli. Attributes included in each image are marked with 1, and those not included are marked with 0. Each attribute has a difference in the number of positive and negative samples.

**FIG 1.**
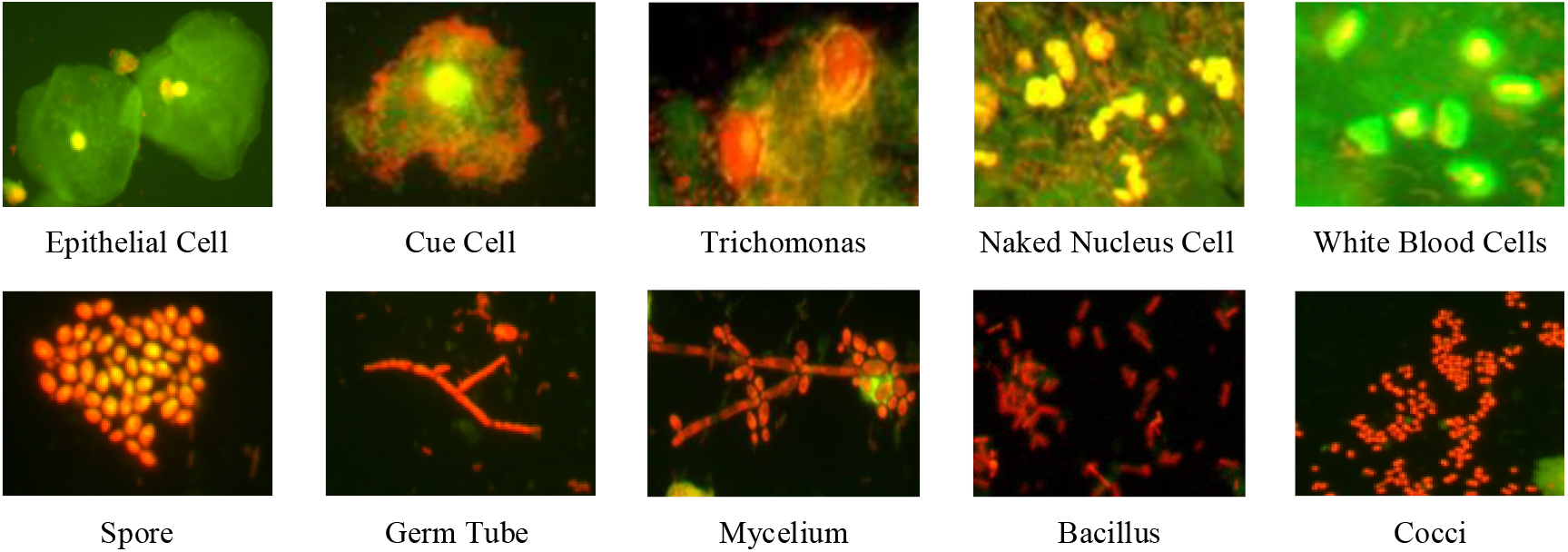
Major bacterial types including inflammatory bacteria and regular colonies in our VB9 dataset.

**FIG 2.**
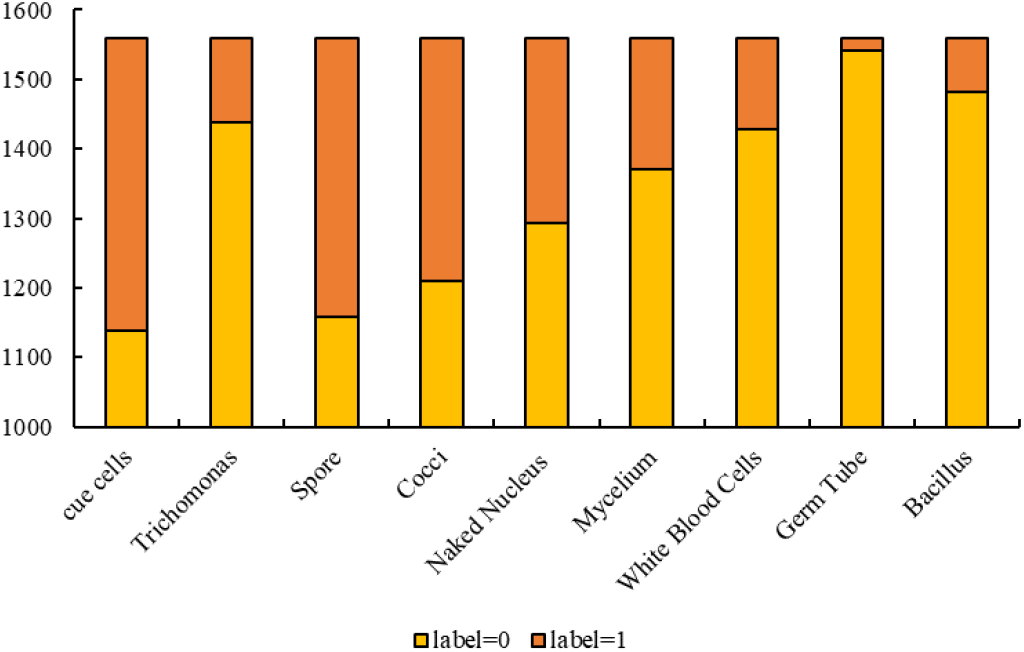
Annotation of different classes of bacteria image attribute in our Vaginitis Bacteria dataset

The method for establishing and processing a multi-label attribute dataset of vaginitis bacteria includes 2 steps. First obtain the bacterial image data, and label the binary one-hot code attribute list according to the bacterial characteristics. Then extract the image and label lists separately, traverse and proofread the image set and attributes so that they correspond one-to-one during training.

### 3.2 Data Augmentation

Deep learning applications rely on good datasets, however annotation of medical data is expensive and cumbersome as it requires a lot of time and expertise from experienced clinicians, making it challenging to annotate large numbers of images. Another major issue is the imbalance of data, which is very common in the health sector. In the field of biomedical imaging, Hao et al.[23] applied data augmentation strategies for prostate cancer detection. To expand the original dataset, we explore several effective data augmentation methods. In general, training data does not provide the location of the bacteria object, so we perform Random Erasing (RE) [24] on the whole dataset. Random Erasing randomly selected a rectangle region *I*_*e*_in an image and erases its pixels with random values, generating training images with various levels of occlusion, the pixels of erased region re-assigned with random values can also be viewed as adding noise to the image.

In training, Random Erasing is conducted with a certain probability. For an image *I* in a mini-batch, the probability of it undergoing Random Erasing and being kept unchanged is *p*and 1−*p*respectively. The area of the training image is *S* = *W* × *H*, the area of erasing rectangle region was randomly initialized to *S*_*e*_, *r*_*e*_is the aspect ratio of erasing rectangle region.

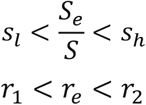

*P*= (*x*_*e*_,*y*_*e*_) is a randomly initialized point in image *I*, when not exceeding the boundary, the selected rectangle area is *I*_*e*_, each pixel in *I*_*e*_is assigned to a random value in [0, 255], respectively.

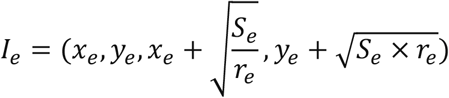

Random cropping and rotation are also effective data augmentation approach, they reduce the contribution of the background, can decrease data noise and increase model stability to a certain degree, in this way, learning models can be built on the existence of object parts rather than focusing on the whole object. Random Image Cropping and Patching (RICAP) [25] randomly crops four images and patches them to construct a new training image, enriching the variety of training images.

We apply the random cropping method, and on this basis add the rotation, flip and affine transformation of the cropped image, random padding of edges is possible when the cropping box exceeds the original image area or when the margin is cropped. Fig 3,4,5 shows examples of rotation and cropping-based augmentation of bacterial image data.

**FIG 3.**
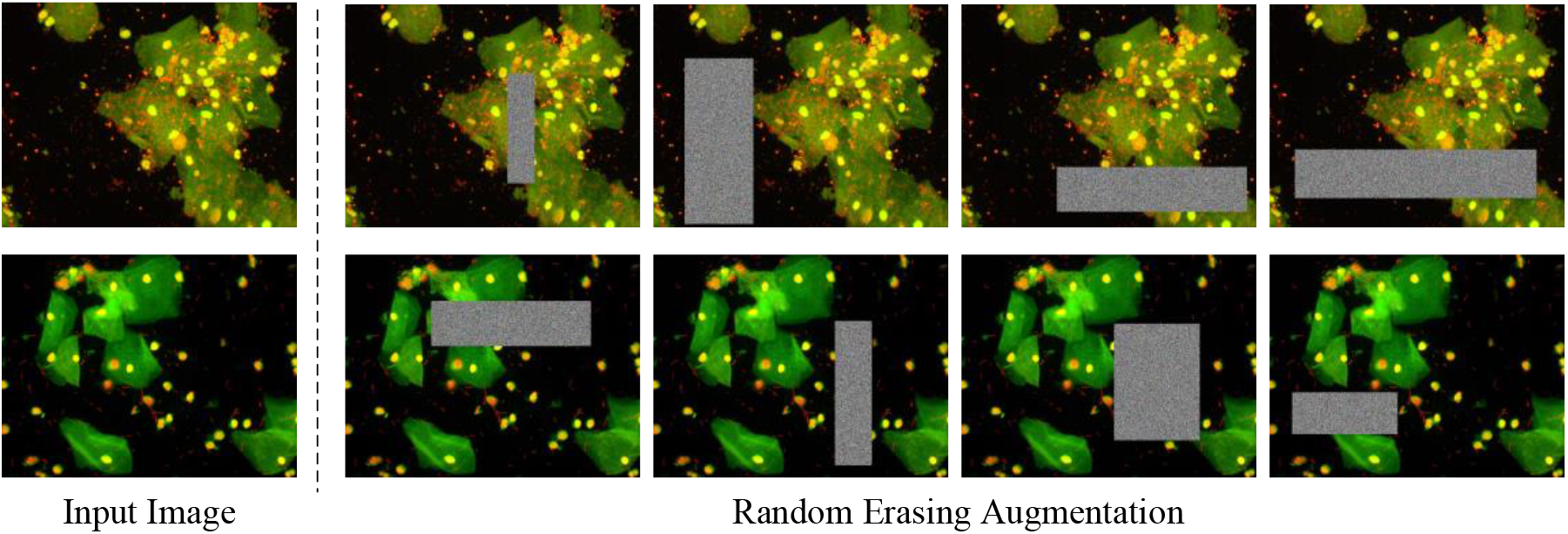
Random Erasing Data Augmentation method, images with various levels of occlusion are generated.

**FIG 4.**
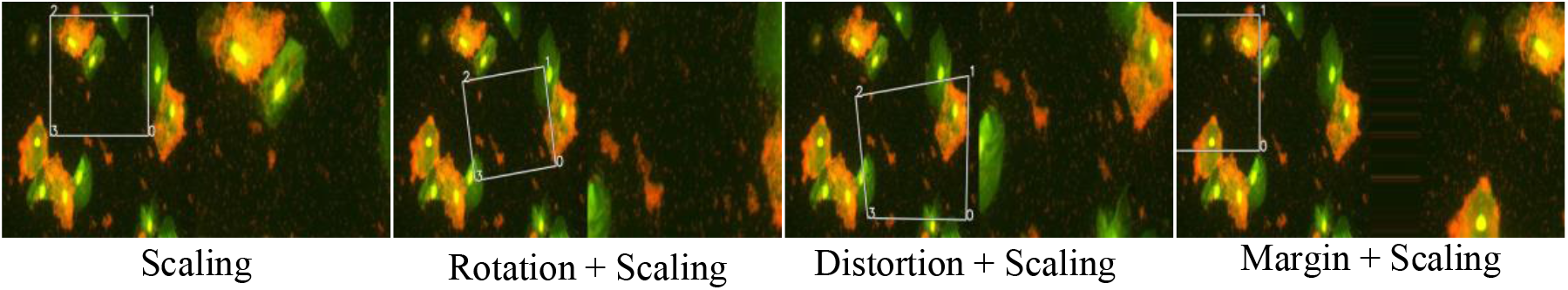
Data Augmentation Method for Random Crop, Rotate, Distort and Edge Filling

**FIG 5.**
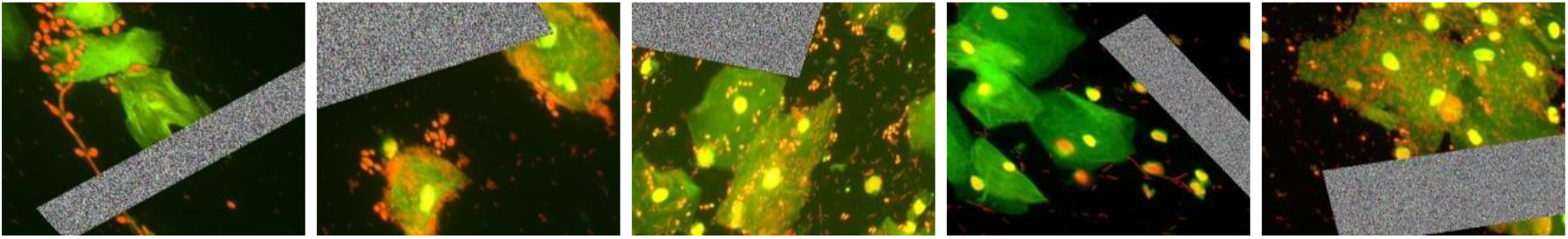
Example of data expansion with random erasure combined with crop rotation

## 4 MODELS AND METHODS

### 4.1 Fine-tuning and deployment of deep models

Convolutional neural networks enable visual patterns to be identified directly from pixel images with minimal preprocessing. Filter number, window size, and convolutional layer depth have been research directions for scholars to improve network performance for many years. On top of the basic network, many derivative networks with excellent performance have also been created. ResNet [16][26] won the 1st place on the ILSVRC 2015 classification task. The skip connection approach allows residual model with a depth of 152 layers still have lower complexity although it is 8 times deeper than VGG [15] networks, which also performed well in classification tasks in ILSVRC Competition in 2014.

There are three main types of related networks for VGG, in addition to VGG (11, 16, 19), there are simplified version SmallerVGG with 5 convolutional layers, and MiniVGG with only 4 convolutional layers in depth. In applications such as multi-label classification or when faced classes with few distinguishing features, experiments [27] showed that lightweight structure of the SmallerVGG avoids the overfitting problem of deep networks. L.J. et al. [27] applied SmllerVGG model on Fine-grained classification task for rowing teams. Murti [28] improved accuracy of identifying isolated Balinese palm leaf characters using SmallerVGG, Allehaibi.et al. [29] applied smaller VGG-like Net on classification of cervical cancer cell segmentation phase and obtained excellent evaluation indicators.

As Fig6 shows, the proposed model uses small filters (3×3 convolution kernels) and applies max pooling layers (2×2) to reduce parameters. The number of multi -label attributes to be identified is equal to output nodes of the end fully connected layer, which uses *sigmoid* activation function as the classifier. In shallow network with fewer convolutional layers, inputting an image with large size will lead to a sharp increase in the number of parameters of the fully connected layer and occupy more computing resources. Images are resized to a suitable dimension (192, 96) before being fed into the model, allowing our network to extract the most effective features.

**FIG 6.**
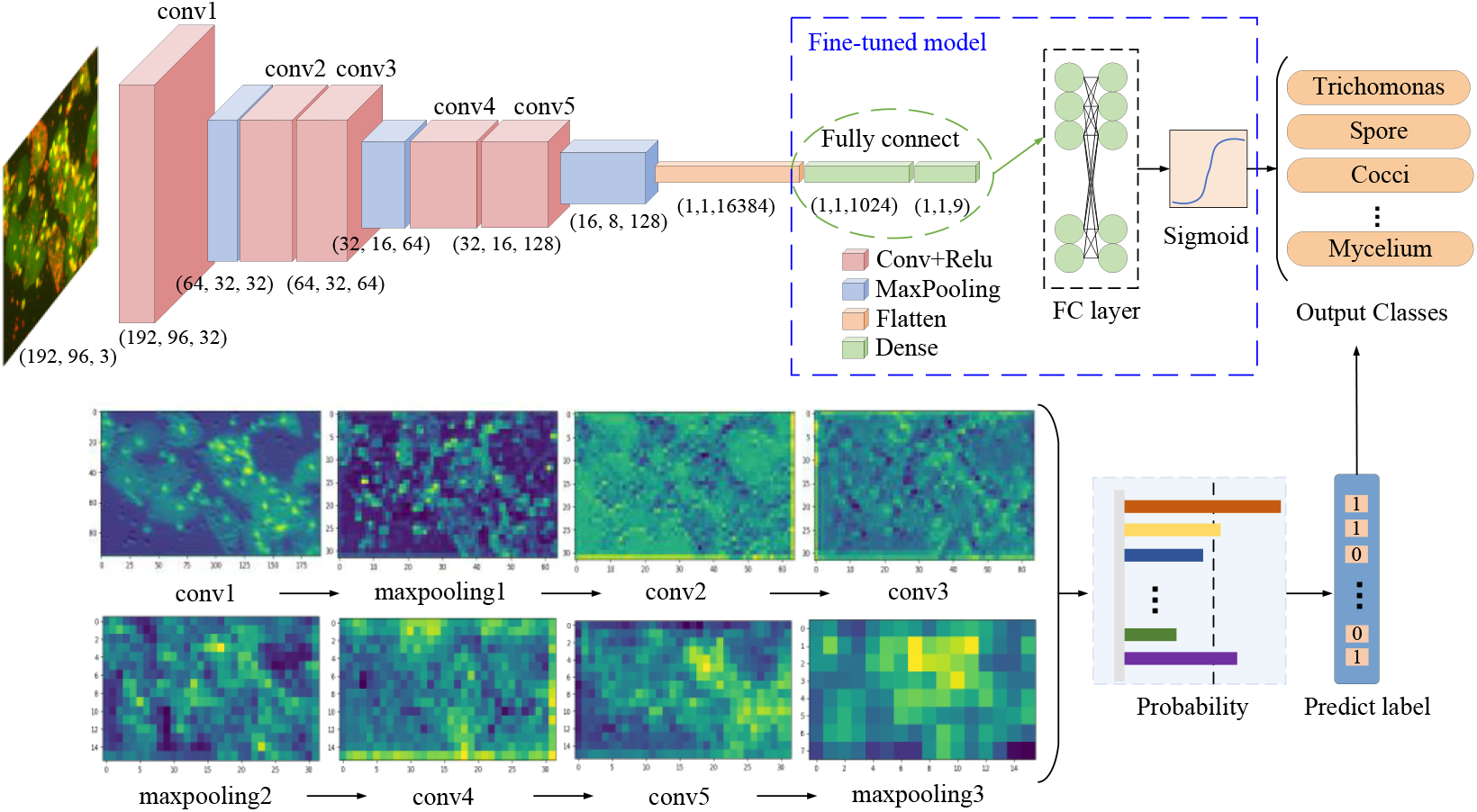
Fine-tuned SmallerVGG (FTS-VGG) model for multi-label classification of vaginitis bacteria with individual convolutional layer visualization and prediction mechanisms

The core of this work is that we have made two improvements on the application of the SmallerVGG model, which makes the proposed model not only lighter and faster, but also shows excellent efficiency on multi-classification tasks, experiments and proofs of which will be given in Chapter 5, the focus of the proposed scheme is:

- ***Fine-tuned end structure:*** We reset fully connected layers for classification with 9 categories, and choose the activation function to be *sigmoid* rather than *softmax*, labelling each output as an independent Bernoulli distribution and penalizing each output node individually.
- ***Binarized unique hot code labels:*** We use one-hot label code and transform the output probabilities of the model into a representation containing only 0 and 1, the presence of attributes is binarized, which allows the calculation for discrete features more reasonable.

To demonstrate the effectiveness of the deployed model, we choose several other basic deep learning backbone networks, including ResNet50, VGG19, InceptionResNetV1, and modify the end structure to apply in this work.

In the field of image classification, transfer learning uses pre-trained models on large labelled datasets such as ImageNet and then fine-tuning on the target dataset, which allows the learned knowledge or patterns to be applied to problems in other domains and effectively accelerates network convergence without compromising model performance. As shown in the Fig7, VGG19 contains a (2, 2, 4, 4, 4) convolutional layer structure with three fully connected layers at the end. It uses successive 3×3 convolutional kernels instead of the larger convolutional kernels in AlexNet, boosting the depth of the network while ensuring the same perceptual field. In this work, we use the output of the last fully-connected layer as image features, add a 9-node fully-connected layer at the end for cell classification, and fine-tune with a pre-trained model without the head for transfer learning.

**FIG 7.**
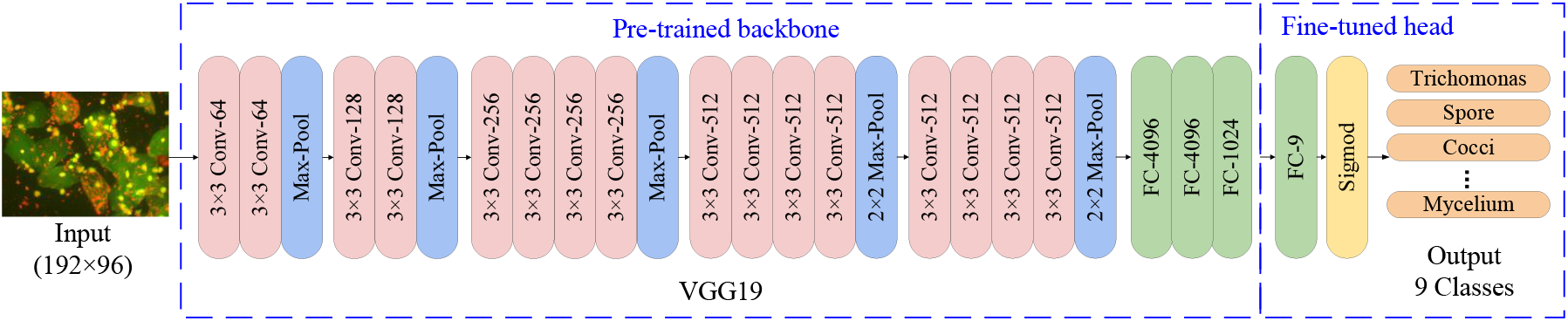
Fine-tuned network architecture with transfer learning on pre-trained VGG19 for multi-label classification of vaginitis bacteria

Fig8 shows the flow of the ResNet50 network applied to this work and the specific structure of Bottleneck. The target cell image input goes through 5 stages of ResNet50 to get the output which is the classification category. Stage 0 preprocesses the input and Stages 1-4 consist of Bottleneck with the structure (3, 4, 6, 3). Bottleneck2 has the same number of input and output channels while Bottleneck1 is different.

**FIG 8.**
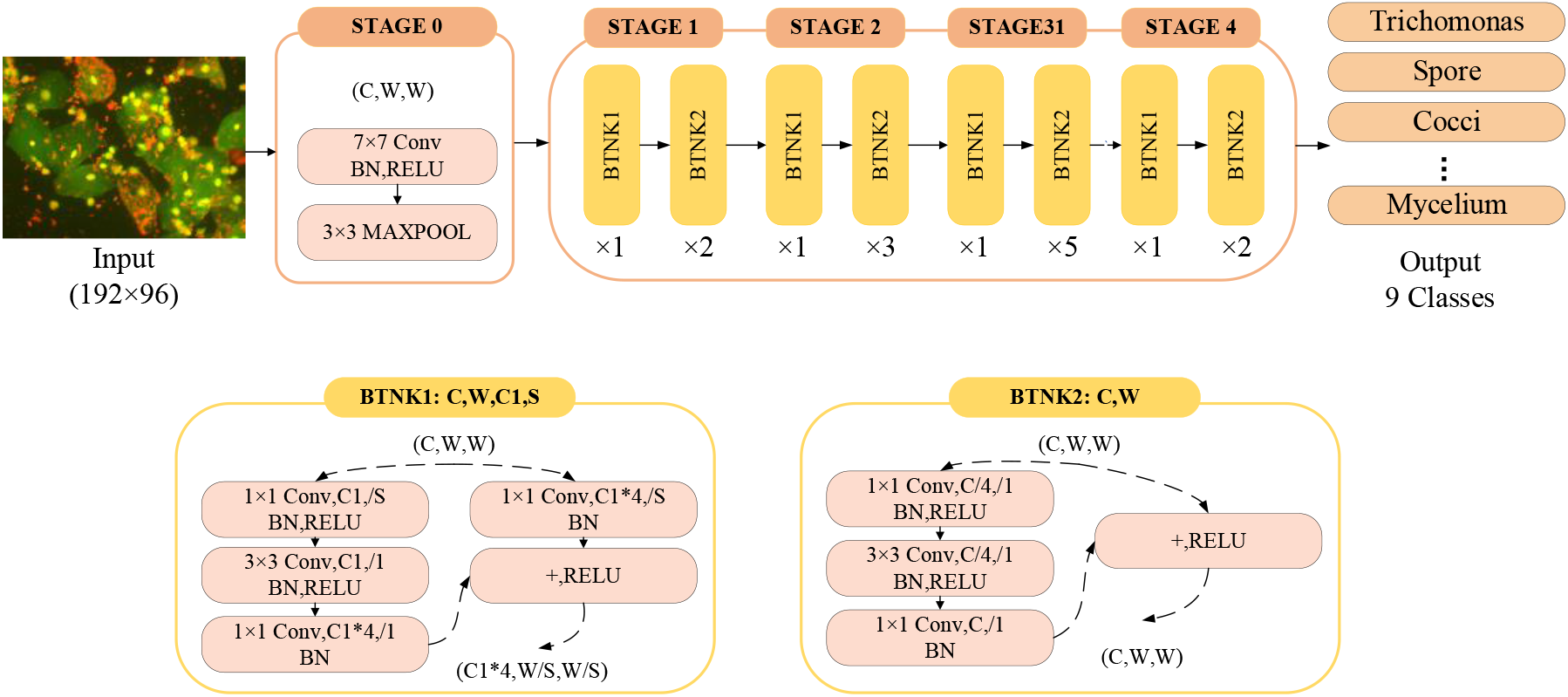
ResNet50 deep network model for multi-label classification

GoogLeNet [12] with Inception module had achieved the best classification and detection performance in ILSVRC14. Inspired by the superior performance of ResNet, researchers proposed a hybrid inception module that introduces residual connectivity, adding the convolutional output of the inception module to the input. InceptionResNet [30] consists of six sub-modules connected successively, including stem, InceptionResNetA, Reduction-A, InceptionResNetB, Reduction-B, InceptionResNetC. As shown in Fig9, in this work we choose the InceptionResNetV1 network, adding a 9-node fully connected layer, a dropout layer and a normalization layer at the end, and finally connecting a sigmoid activation layer for classification.

**FIG 9.**
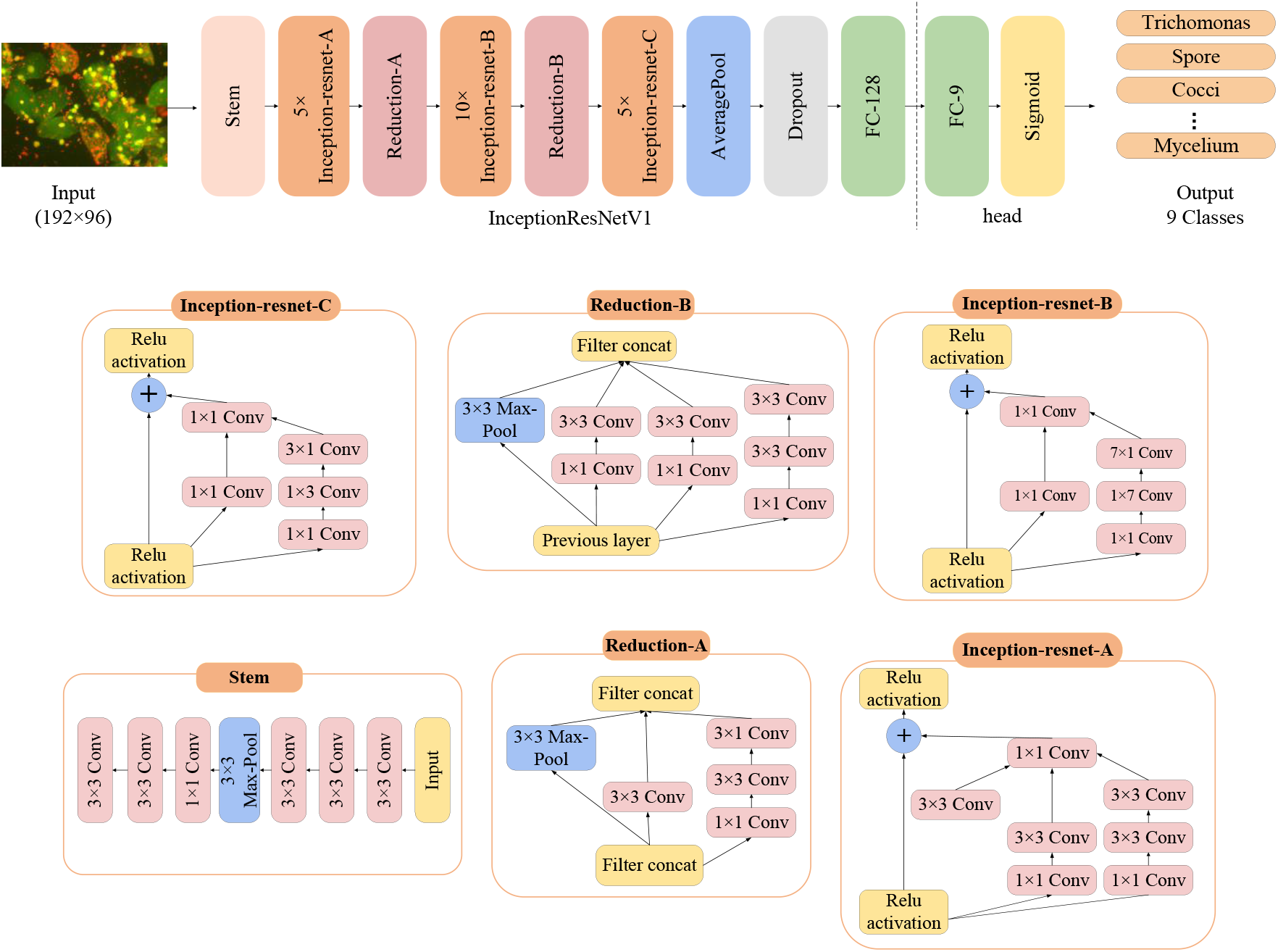
InceptionResNetV1 deep network model for multi-label classification

### 4.2 Loss Function

In bacterial classification task, the proposed model obtains the probability value of each predicted attribute through *sigmoid* calculation. During the training process, the performance of the model on the sample is judged by the loss function, which is continuously optimized to improve the network performance.

In the case of dichotomous categories that distinguish between negative and positive, the model ultimately predicts only two types, with each category predicted with probabilities of *p*and 1−*p*.

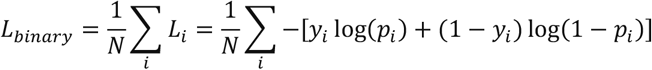

*y*_*i*_ represents the label of the sample, the positive class is 1, and the negative is 0. *p*_*i*_ represents the probability that the sample is predicted to be a positive class.

Based on the simple classification, the multi-label classification proposed in this work simultaneously predicts multiple bacterial attributes with a loss function further expressed as:

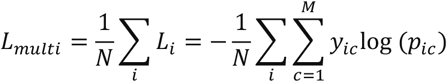

*M* is the number of categories, *y*_*ic*_is the sign function that takes 1 if the true category of sample, *i* is equal to *c* and 0 otherwise, *p*_*ic*_is the predicted probability that the observed sample *i* belongs to category *c*.

### 4.3 Evaluation Metrics

For multi-label bacterial classification and recognition, average precision (mAP) is generally used as an evaluation metric, which reflects the property of ranking the correct search results at the top of the test list. The average precision (AP) is the area under the PR (Precision-Recall) curve, where the horizontal and vertical axes are usually Recall and Precision, respectively. In addition, the closer the PR curve is to the upper right corner, the better the model performance is. Precision and recall are calculated as follow :

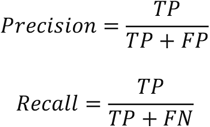

where TP (True Positives) is the number of detected frames with IOU > 0.5; FP (False Positives) indicates the number of redundant detected frames with the same true label; TP, FP, TN and FN indicate true positive, false positive, true negative, and false negative, respectively. Further, recall is also referred to as the hit rate or true positive rate (TPR), correspondingly, the false positive rate (FPR) and the true negative rate (TNR) are defined:

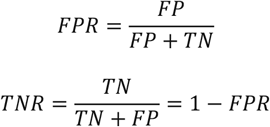

F1 score is the harmonic mean of the precision and recall, which takes both false positive samples and false negative samples into account:

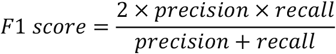

Identification results for each bacterial attribute are evaluated by accuracy, which is the ratio of correct predictions to the total number of predictions, the calculation formula is given by:

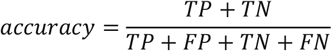

The Receiver Operating Characteristic (ROC) curve takes FP and TP as horizontal and vertical coordinates respectively, showing the trade-off between sensitivity and specificity, which can visually represent the performance of the classifier. The area of the part under ROC curve is defined as AUC, and the value is usually between 0.5 and 1.0, with higher AUC representing better performance.

Confusion matrix counts the number of misclassified and right-classified observations of the model, and summarizes the records in the dataset by true and predicted category in a matrix form, which are expressed as the rows and columns, respectively.

## 5 EXPERIMENTS AND RESULTS

We firstly employ SmallerVGG network as the basis for classification stage. The images after the convolution and pooling are flattened and fully connected to 1024 dimensions by a dense layer as image features, followed by the end fully connected layer, whose number of output nodes is equal to the bacterial attribute category. Sigmoid is selected as the activation function at the end of the model to non-linearize each output value of the network, the scalar number is converted to [0, 1] for binary classification to determine whether the attribute exists. The Adam optimization method and the binary cross-entropy loss were chosen because the multi-label classification treats each predicted label as an independent Bernoulli distribution to evaluate each output node individually.

During training phase, the RGB pixel values in the [0,255] are normalized in the pre-processing stage to obtain absolute color information, avoiding the interference caused by unbalanced pixel luminance distribution while ensuring faster convergence. According to the order of image loading, image indices are used to rearrange the label list, so that the real value of the attribute corresponds to the image one-to-one. Then images are randomly divided into training set and test set and shuffled to make sure that the arrangement of data has a certain randomness and the possibility of any type of data is the same when reading sequentially.

In the testing and evaluation stage, first load the trained model and ground truth labels, then process the image to be tested and input into the net. The attribute index is marked according to the output and threshold value, which is used to judge the prediction of the bacterial species type. To quantitatively evaluate the test performance, we calculate the accuracy, precision and recall at the image level and attribute level respectively, finally construct a dictionary of background and prediction results to print the attribute identification and evaluation results. Specifically, for each image level, the predicted value *i*_*pre*_of length 9 and the true label *i*_*truth*_are used to calculate the evaluation metrics. For various attribute levels, we obtain the attribute predictions *n*_*pre*_and true label *n*_*truth*_of length *n* by splicing *i*_*pre*_and *i*_*truth*_of n images respectively.

In order to visualize the efficiency of the test model, we use key-value pairs to organize the attribute prediction results, where the keys are 9 attribute categories, and the values are the output of the net, representing the probability of each type of bacteria in all possible kinds.

### 5.1 Comparison of Original and Augmented Datasets

We perform cross-training and validation on the raw data and enhanced images respectively, exploring continuously to improve the accuracy of the model. Due to the limitations of various factors, we collected 100 raw data initially, and obtained 83.11% accuracy and 54.93% precision for training. After expanding 50 pieces of data, the accuracy and precision are increased to 92.37% and 74.5%, respectively. As shown in Fig11, then we augment the data with random erasure, rotation, and cropping as described in Section 3.2, retrain the model and validate it on raw data, cropped data, REA data, and all augmented data. Fig12 shows results from training and testing on 6 different datasets, with a significant improvement in evaluation metrics on the enhanced dataset. Table1 corresponds to Fig12 and gives specific values for accuracy, precision and recall on 6 datasets, also reflects the process of improving the performance of our model step by step through data enhancement. It can be seen that the accuracy fluctuates above 97.42%, and the result obtained on the added image achieves the best performance of 98.67%. In addition, the training process shows that the original small dataset takes 200 epochs or even more time for network fitting, while the enhanced dataset only uses about 50 epochs to converge.

**FIG 10.**
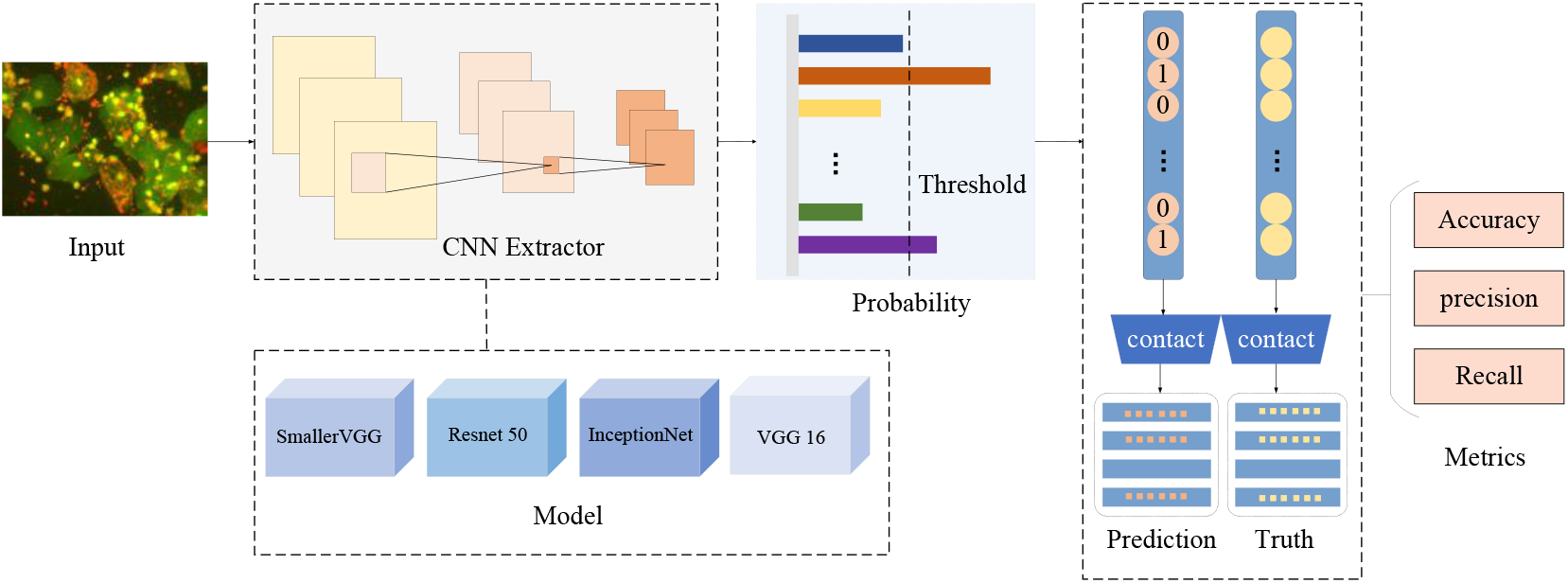
The proposed multi-label classification model workflow and evaluation methodology for vaginosis bacterial

**FIG 11.**
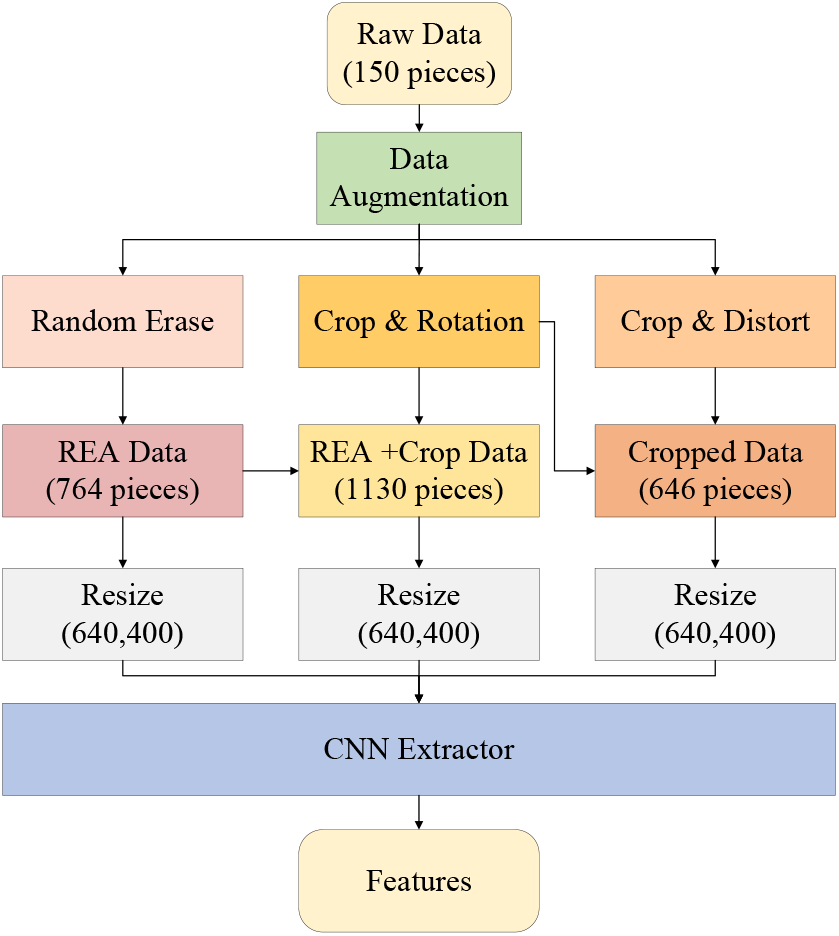
Composition and expansion of the dataset

**FIG 12.**
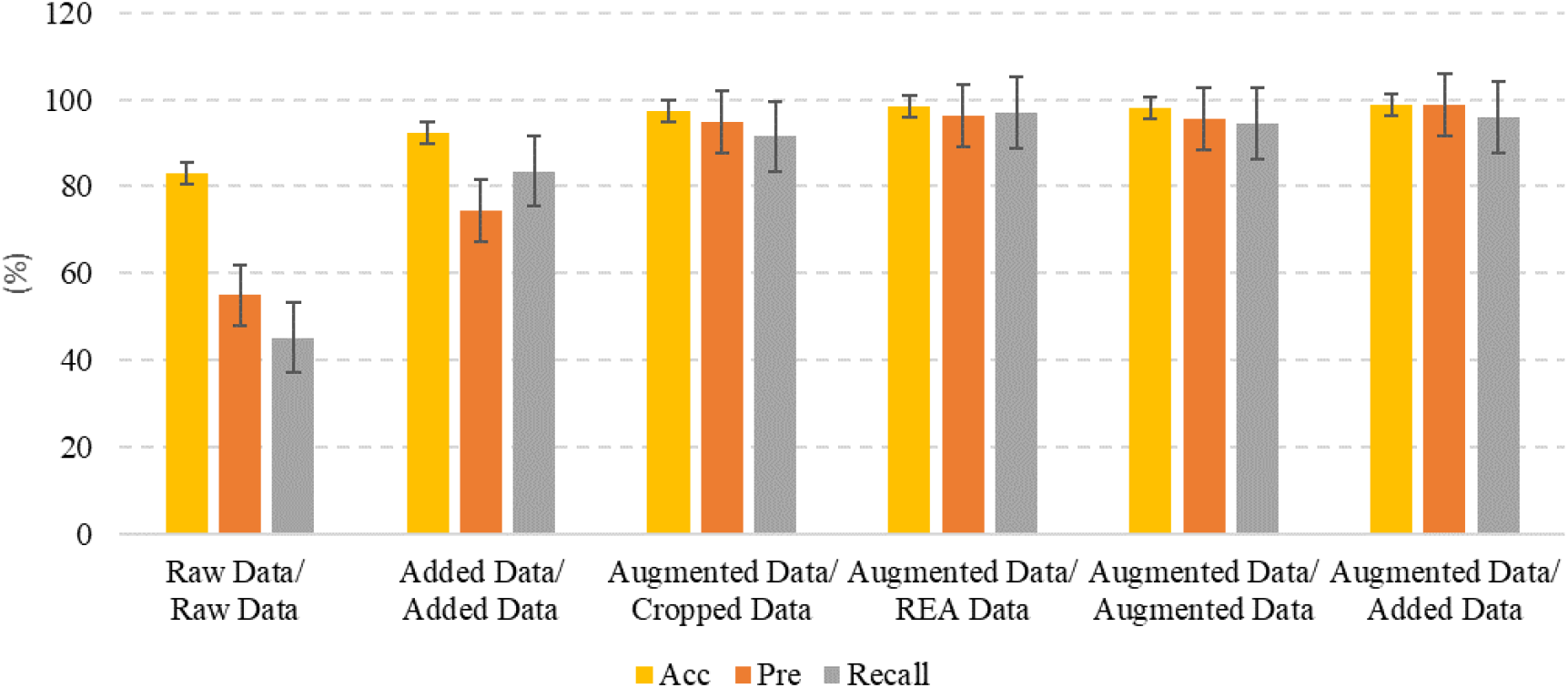
Performance improvement of the proposed model during data expansion

**FIG 13.**
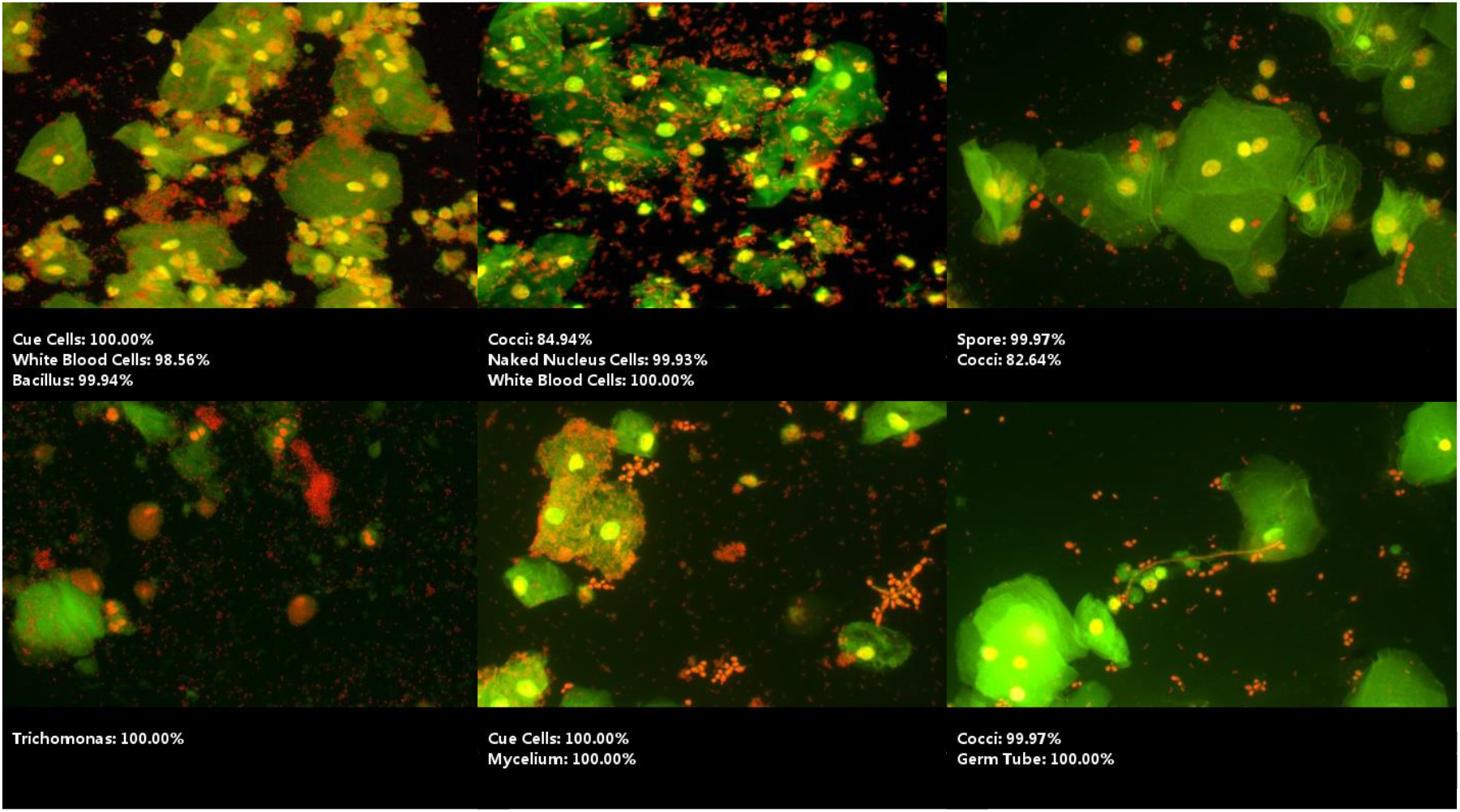
Sample results of multiclassification bacteria identification based on the proposed Fine-tuned SmallerVGG model

**TABLE 1.** Comparison of evaluation metrics on different test sets and train sets

| Traindata | Raw Data | Added Data | Augmented Data |  |  |  |
| --- | --- | --- | --- | --- | --- | --- |
| Testdata | Raw Data | Added Data | Raw Data | REA Data | Crop Data | Augmented Data |
| Acc | 83.11% | 92.37% | 98.67% | 98.60% | 97.42% | 98.05% |
| Pre | 54.93% | 74.50% | 98.68% | 96.38% | 94.74% | 95.53% |
| Recall | 45.15% | 83.53% | 95.97% | 97.08% | 91.49% | 94.46% |

Fig14 shows the confusion matrix for our best performing model tested on augmented dataset, each bacterial type can be considered as an attribute, the diagonal line of each matrix plot shows the correct result for identifying a negative positive for specific type.

**FIG 14.**
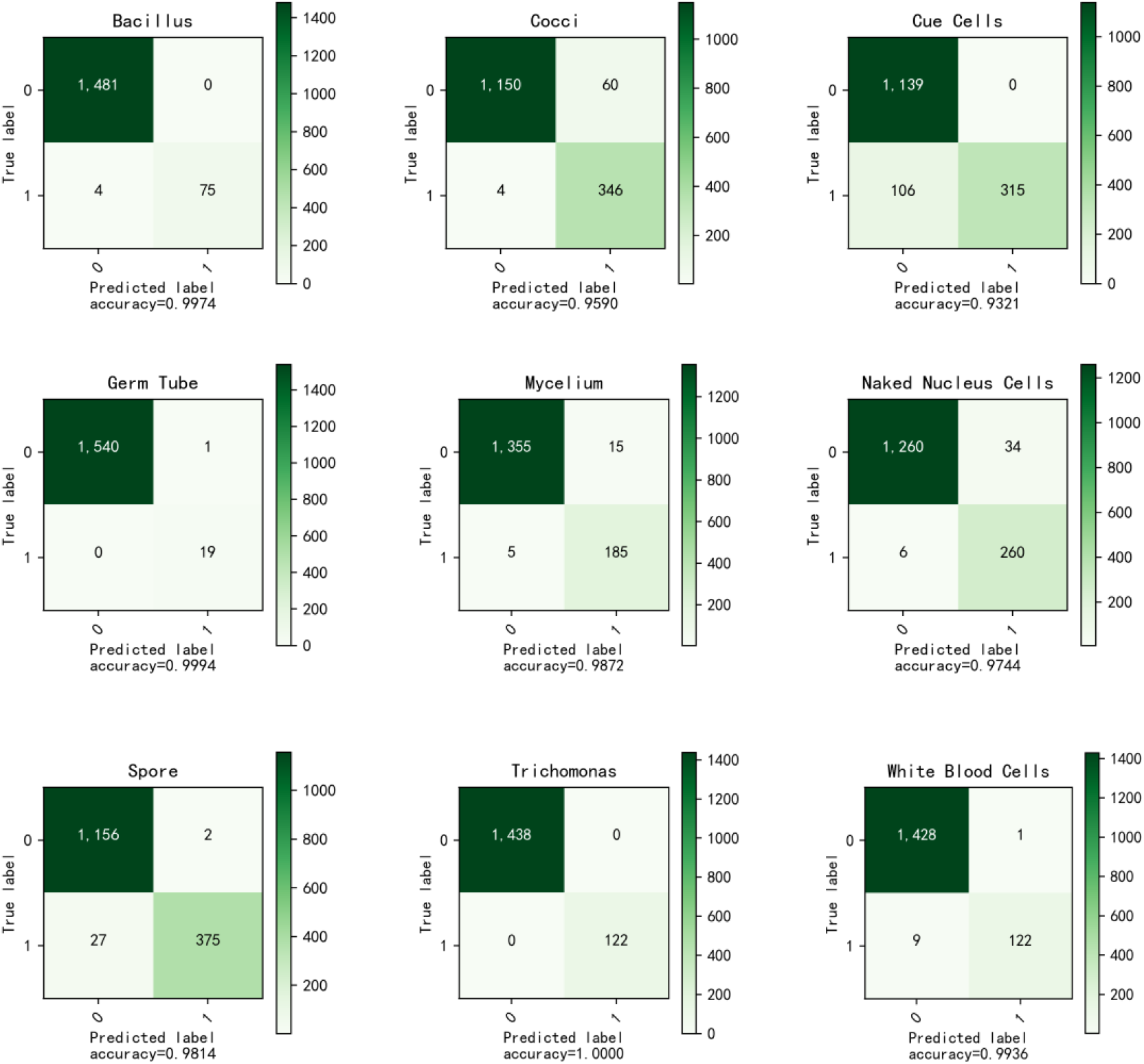
Confusion matrix for multi-label classification of 9 bacterial types tested on augmented dataset by the proposed model

### 5.2 Comparison of Efficiency of Different Backbone Models

#### 5.2.1 Model construction and parameters

In section 4.1 we discussed the structure of the 4 types deep learning models and described how they are deployed for multi-label classification tasks. Table2 shows the network parameters of these models. Fine-tuned SmallerVGG with 5 convolutional layers has the fewest parameters, while the 3 convolutional layers at the end of VGG19 and the 9-node convolutional layer for classification lead to a dramatic increase in the number of parameters. ResNet and InceptionResNetV1 networks have a deeper structure with 10 million more parameters than SmallerVGG, consuming huge computational resources. In comparison, our FTS-VGG for multi-tag classification is the lightest and easier to deploy in practice.

**TABLE 2.** Network parameters for different deep learning models

| Parameters | InceptionResNetV1 | ResNet50 | VGG19 | FTS-VGG |
| --- | --- | --- | --- | --- |
| Train time | 4480 | 7623 | 14498 | 1776 |
| Total params | 22,594,539 | 23,590,793 | 29,475,913 | 17,070,985 |
| Non-trained params | 28,594 | 45,440 | 2,048 | 2,880 |

#### 5.2.2 Training processes and convergence rate

We recorded the training details of the 4 models during 100 epochs. Fig15 shows the changes in loss and accuracy on the enhanced dataset. It can be seen that, first, Fine-tuned SmallerVGG has reached stable convergence around the 70th generation, while VGG19 and InceptionResNet still show a downward trend in loss. Table 2 shows that FTS-VGG takes the shortest time to train, and VGG19 takes almost 8 times as long, which indicates that our model is faster. Second, although the InceptionResNetV1 network is deeper and has more parameters, and no pre-trained model was used but started from scratch, the trend of convergence is generally the same as that of VGG. One detail is that compared with VGG, it uses much less time per epoch, this is because of the effect of the residual block, which is also illustrated by the rapid convergence of ResNet.

**FIG 15.**
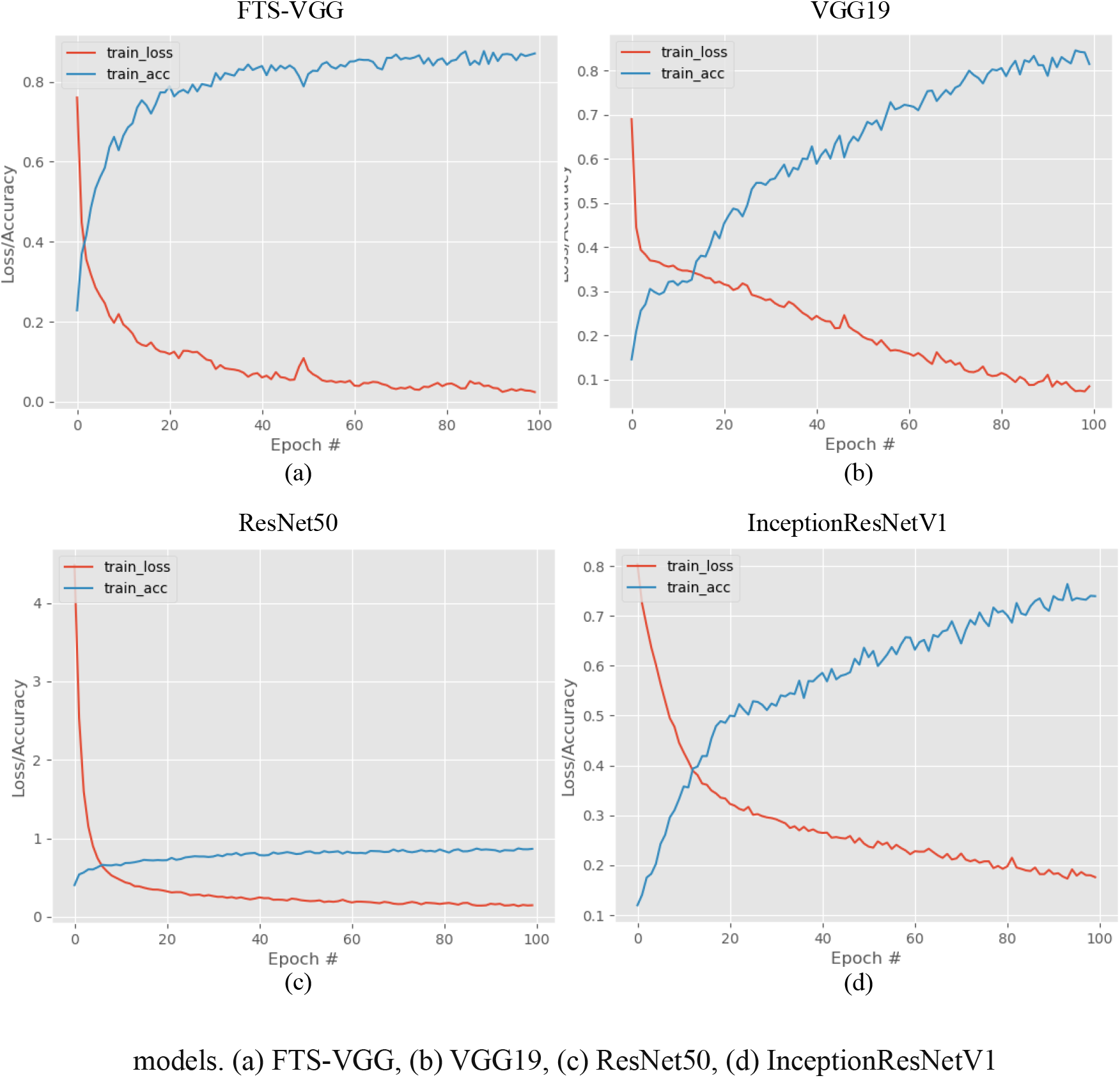
Training process and convergence rate on augmented dataset of several deep learning

By comparison we conclude that our proposed model based multi-classification network has the shortest training time and the fastest convergence rate. In addition, we give clear empirical evidence that the application of residual blocks can significantly speed up the training of the ResNet and InceptionResNet networks.

#### 5.2.3 Model efficiency and evaluation metrics

We tested and evaluated the four models trained on the augmented dataset as described in Section 5.2.2. Fig16 shows a comparison of their performance metrics, with FTS-VGG achieving 98.96% accuracy, 96.79% precision and 98.65% recall, outperforming the other applied backbone networks. As can be seen from the specific values in Table 3, the accuracy rates of the four models are all over 94%, and the recall rate and precision rate are quite different. Accuracy of IncpetionResNetV1 is second only to the best performing FTS-VGG, but the other two metrics are lower.

**FIG 16.**
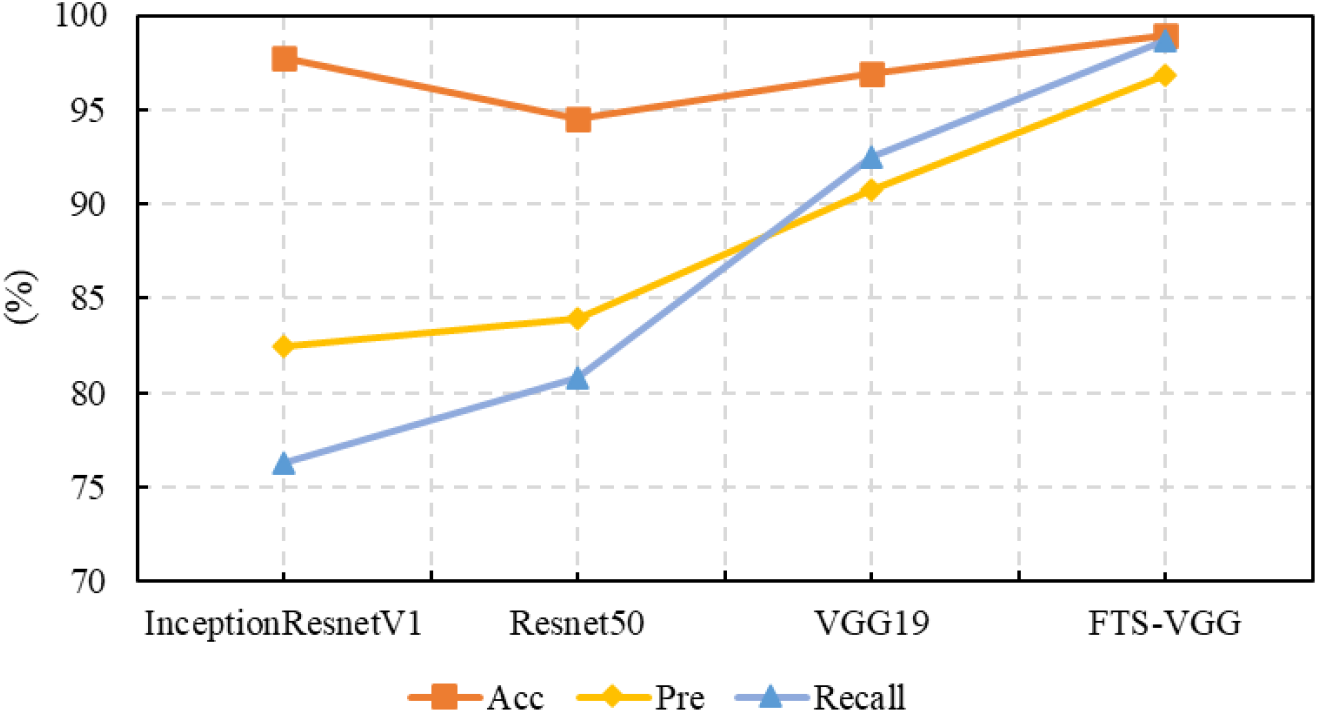
Comparison of the accuracy, precision and recall of the models applied in this work

**TABLE 3.** Network parameters for different deep learning models

| Metrics | InceptionResNetV1 | ResNet50 | VGG19 | FTS-VGG |
| --- | --- | --- | --- | --- |
| Acc | 97.78% | 94.52% | 96.89% | **98.96%** |
| Pre | 82.45% | 83.89% | 90.74% | **96.79%** |
| Recall | 76.31% | 80.78% | 92.49% | **98.65%** |

Fig17 shows the ROC curves for 4 comparative models in a multi-classification test for 9 bacterial type attributes. The curves use the false positive rate (FPR) and true positive rate (TPR) as the horizontal and vertical coordinates, respectively, the closer their area is to 1, the better the recognition performance, with complete recognition indicated when the area equals 1. Specifically, the area AUC value below the curve corresponding to each cell is marked in the bottom right corner of each graph. The advantage of ROC curve is that it is very intuitive to see that the 9 curves of FTS-VGG model are closest to the top of the axes, implying a larger AUC value and indicating the most superior performance of our chosen model.

**FIG 17.**
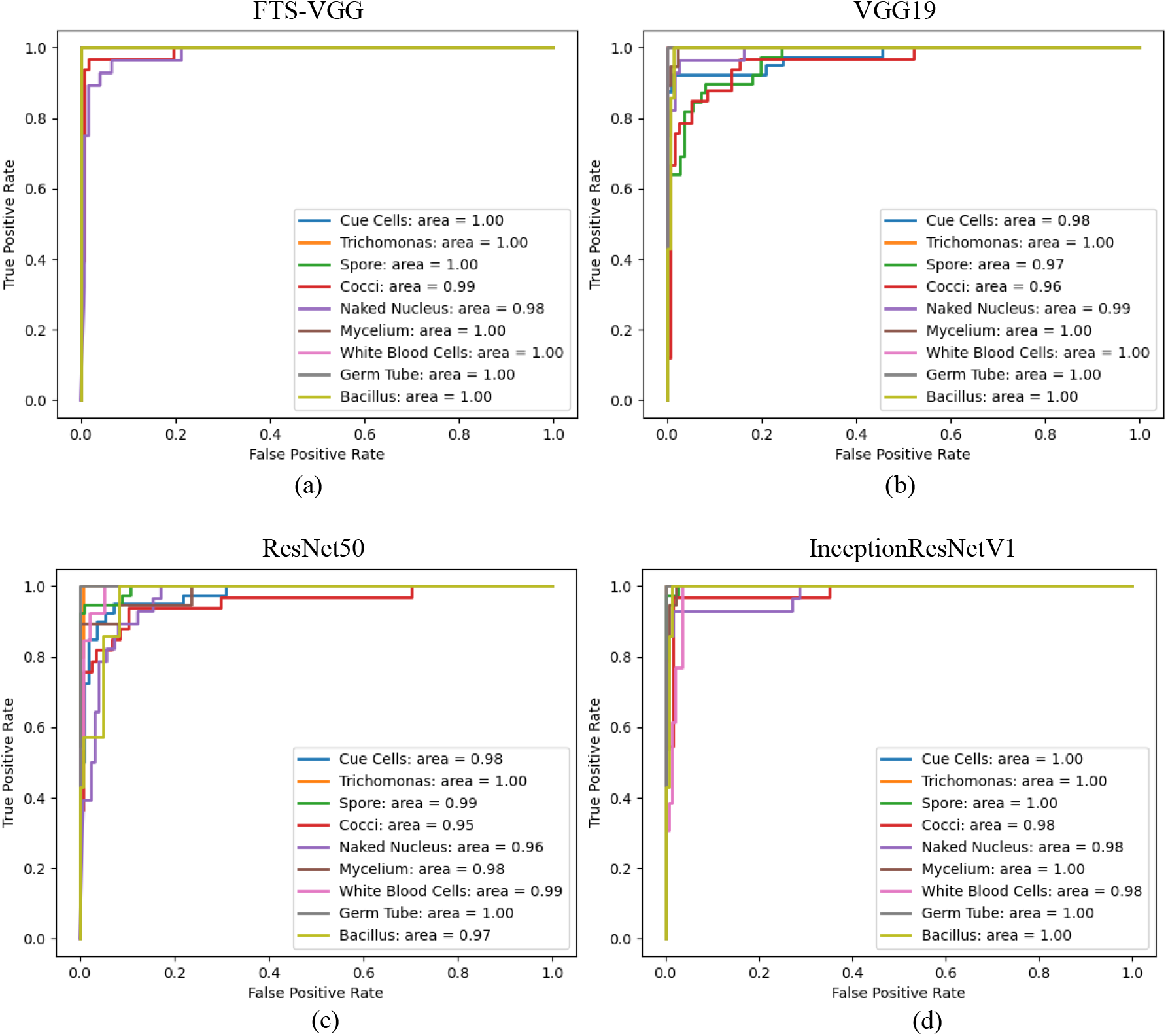
ROC curves with AUC values of 4 deep learning models for multi-classification test for 9 cell attributes. (a) FTS-VGG, (b) VGG19, (c) ResNet50, (d) Inception ResNetV1

The comparisons in this subsection lead to the conclusion that the model proposed in this work performs best in terms of each evaluation metric and is largely adequate in practical applications.

## 6 CONCLUSION AND FUTURE WORK

In this paper, we proposed a Fine-tuned SmallerVGG based multi-label identification method for vaginitis bacteria, obtained highest test accuracy of 98.67% and best AUC results for 9-class classification of Deep CNNs. When multiple different species of bacteria appear simultaneously, they are regarded as image attributes. Deep convolutional networks are used to extract features and output predicted probabilities. We apply random erasure, rotation and cropping for data augmentation to solve the problem of small datasets and gain high accuracy. Marriage of high-performance computing with machine learning promise the capacity to deal big medical image data for accurate and efficient diagnosis. Experiments were performed on 4 advanced backbones and then tested and evaluated at the image level and attribute level, respectively. Analyzing the experimental results, we can draw the following conclusions:

- Data Augmentation methods can further improve the performance of bacterial recognition while accelerating the convergence of model training.
- Higher validation on REA data indicates that Random Erasing can makes the model robust to occlusion and reduce the risk of over-fitting.
- Comparison of model structure and parameters, convergence speed and evaluation

metrics strongly suggest that the proposed fine-tuned model is lighter and faster, more accurate and more efficient.

In the future we will continue to build on this foundation and develop new models to enable location identification and segmentation. The model presented in this work can be further modified to diagnose more types of cells. It is expected to develop into a core method for automated diagnostic equipment that is more accurate, efficient and stable than manual diagnosis.

## ACKNOWLEGMENTS

This work is supported by National Natural Science Foundation of China (51975360, 52035007); National Social Science Foundation of China (17ZDA020); Cross Fund for medical and Engineering of Shanghai Jiao Tong University (YG2021QN118).

